# Identification of theTMEM232 gene associated with atopic dermatitis through targeted capture sequencing

**DOI:** 10.1101/2021.02.25.432973

**Authors:** Jie Zheng, Yuan-yuan Wu, Wen-liang Fang, Xin-ying Cai, Zeng-yun-ou Zhang, Chong-xian Yu, Xiao-dong Zheng, Feng-li Xiao

## Abstract

**BACKGROUND:** Atopic dermatitis (AD) is a common and complex skin disorder, and the 5q22.1 region had been reported to be associated with AD.

**OBJECTIVE:** To identify the susceptibility gene for AD in the 5q22.1 region by haplotype and targeted capture sequencing.

**METHODS:** The haplotypes were reconstructed with our previously genotyping data of four SNPs and six deletions from 3055 Chinese Hans AD patients and 4346 controls. The targeted capture sequencing spanning 5q22.1 region was performed in the selected samples. The gene level enrichment analysis was done using loss of function variants.

**RESULTS:** A total of 60 haplotypes were found, and the H15 haplotype had the strongest association with AD (*P* = 2.72×10 ^−10^, OR = 0.14, 95%CI = 0.07-0.28). However, No co-segregation mutation sites were found in the sequencing analysis within the 16 selected samples, while the enrichment analysis indicated that TMEM232 was significantly associated with AD (P = 7.33×10 ^-5^, OR = 0.33, 95% CI = 0.19-0.58).

**CONCLUSIONS:** This study confirms previous findings that the TMEM232 gene is associated with AD by haplotype analysis and targeted capture sequencing.

## 1. Introduction

Atopic dermatitis (AD) is a common, multifactorial chronic inflammatory skin disease ^[1]^. In recent years, the number of AD cases has gradually increased with a prevalence of 0.2% to 24.6% worldwide ^[2]^. The aetiology of AD remains unclear. The dysregulation of immune response and defects in the epidermal barrier has a critical impaction in the development of AD, which was caused by the interactions between genetic and environmental factors ^[3, 4]^.

We have conducted a genome-wide association study (GWAS) of AD among the Chinese Han population in 2011, the previously undescribed susceptibility loci at 5q22.1(TMEM232 and SLC25A46) was identified to have an association with AD^[5]^, which was subsequently replicated in a Japanese AD GWAS^[6]^. To identify possible susceptibility genes in 5q22.1, we further scanned the indels and single-nucleotide polymorphisms (SNPs) in the 5q22.1 region through genomic imputation and genotyping. Six deletions and four SNPs were associated with AD. The strongest variant rs11357450 deletion is located in TMEM232. The protein expression of TMEM232 was different between AD and normal tissues by immunohistochemistry. TMEM232 may be a susceptibility gene for AD in the Chinese Han population^[7]^.

The haplotype may be more robust than the individual markers for delineating the susceptibility genes of complex disease ^[8]^. The reconstructed haplotypes analysis were more reliable, cost-effective method to predict an individual’s reaction to a drug or risk of disease rather than single SNPs ^[9]^. The haplotype studies of AD provided a molecular basis for explaining the pathogenesis of disease. Lacy et al^[10]^ found IgE levels in AD patients was associated with the haplotype TGAC in the IL10 promoter region. Tang et al ^[11]^ performed genetic association study of the genotypes and haplotypes and implicated that SHARPIN may be a novel participant in the pathogenesis of AD. Our study intends to define haplotypes from the previous genotyping data in the 5q22.1 region^[7]^. High-throughput sequencing was used to discover whether there was a high-risk haplotype would co-exist with a gene function mutation, and loss of function (LOF) variants enrichment test in gene level will be performed to explore candidate genes for AD.

## 2. Materials and methods

### 2.1. Study samples

This study was performed on 3055 patients (1846 men and 1209 women), with a mean age of 6.87±8.95 years old, and 4346 controls(2256 men and 2090 women) with a mean age of 28.16±13.75 years old. All participants were unrelated and of Chinese Han origin. The clinical information was collected through comprehensive clinical examinations. The diagnosis of AD was made by at least two experienced dermatologists based on the standard criteria of Hanifin and Rajka criteria^[12]^. All control groups were healthy individuals without AD, other atopic diseases, systemic diseases, or a family history of AD (including first-, second- and third-degree relatives). All participants or their guardians received written informed consent. The study was conducted in accordance with the Declaration of Helsinki principles and was approved by the Institutional Ethics Committee of Anhui Medical University.

### 2.2. Defining haplotypes

The genotyping data of the significant four SNPs (rs10067777, rs7701890, rs13360927 and rs13361382) and six deletions(rs5870408,rs140764268,s11357450, rs35639206,rs137936676,rs10617471)were extracted spanning the 5q22.1 region from our previous AD studies^[5, 7]^. These variants were in strong linkage disequilibrium (LD: r2 ≥ 0.80). The PHASE v 2.1 was used for reconstructing haplotypes from our AD genotype data ^[13, 14]^. Then, the local Perl script was applied to convert the polymorphic results of haplotypes into biallelic format, and the p-value and corresponding odds ratio (OR) with a 95% confidence interval were computed using Chi-square tests implemented in PLINK version1.07.

### 2.3. Selecting samples for sequencing

Some samples were selected for sequencing. Firstly, the samples with the top significant haplotypes were prioritized. Secondly, one with this homozygote haplotype was screened as many as possible, and that with heterozygotes haplotype were selected when a sufficient number of homozygotes were not available. Third, the selected samples represented each high frequency haplotype (> 0.5%). At last, two samples without the haplotype were screened as a reference.

### 2.4. Targeted next generation sequencing

The genomic DNA samples that met the criteria were randomly fragmented by Covaris technology. The size of the library fragments was mainly distributed between 150 bp and 250 bp. The end of DNA fragments was repaired and an “A” base was added at the 3’-end of each strand. Adaptors were then connected to both ends of the end repaired/dA tailed DNA fragments for amplification and sequencing. Selected DNA fragments were amplified by ligation-mediated PCR (LM-PCR), purified, and hybridized to the targeted region array for enrichment. Non-hybridized fragments were then removed by washing. Captured products were then circularized. DNA nanoballs (DNBs) were produced by rolling circle amplification (RCA). Each qualified captured library was then loaded on the BGISEQ-500 sequencing platform and high-throughput sequencing is performed on each capture library to ensure that each sample met the required average sequencing coverage. The targets covered approximately 2.9 Mb (from 109M to 112M) of the 5q22.1 region and its adjacent sequences.

### 2.5. Mapping and variants calling

To reduce the noise of sequencing data, data filtering was carried out as follows: removing reads containing sequencing adaptors; removing reads whose low-quality base ratio (base quality less than or equal to 5) is more than 50%; and removing reads whose unknown base (‘N’ base) ratio is more than 10%. Statistical analysis of data and downstream bioinformatics analysis were performed on this filtered, high-quality data, referred to as the “clean data”.

All clean data of each sample were mapped to the human reference genome (GRCh37/HG19). The alignment was performed by Burrows-Wheeler Aligner (BWA) software. To ensure accurate variant calling, we followed the recommended Best Practices for variant analysis with the Genome Analysis Toolkit (GATK, https://www.broadinstitute.org/gatk/guide/best-practices). GATK was used to recalibrate base quality score and realign local around indels, with duplicate reads removed by Picard tools. HaplotypeCaller of GATK was used to detected genomic variations, including SNPs and indels. The variant quality score recalibration (VQSR) method, which uses machine learning to identify annotation profiles of variants that are likely to be real, was used to obtain high-confident variant calls, and all variant calls were annotated by ANNOVER.

### 2.6. Gene level enrichment analysis

Variants annotated by ANNOVA as exonic, UTR, splicing site or upstream will consider to be LOF variants. All these variants were detected manually to find out whether they were co-segregation with the most significant haplotype. With an in-house perl script to count LOF variants number in case and control cohorts respectively, the gene level enrichment analysis was performed using chi-square test on genes that had two or more LOF variants to detect the susceptibility genes of AD.

## 3. Results

### 3.1. The results of haplotype analysis

A total of 60 haplotypes were found in the discovery stage (Supplementary Table 1). The most frequent haplotype was H1, with a frequency of 21.8% in cases and 21.4% in controls (*P* = 0.56, OR = 1.02, 95% CI = 0.95-1.11). A significantly higher frequency of H60 (*P* = 7.39×10 ^−4^, OR = 1.26, 95% CI = 1.10-1.44) was detected in patients than in controls. The frequencies of H15, H14 and H52 were obviously lower in patients than in controls. And the most significant association was observed between H15 and AD (*P* = 2.72×10 ^−10^, OR = 0.14, 95% CI = 0.07-0.28) (Table 1).

**Table 1.**
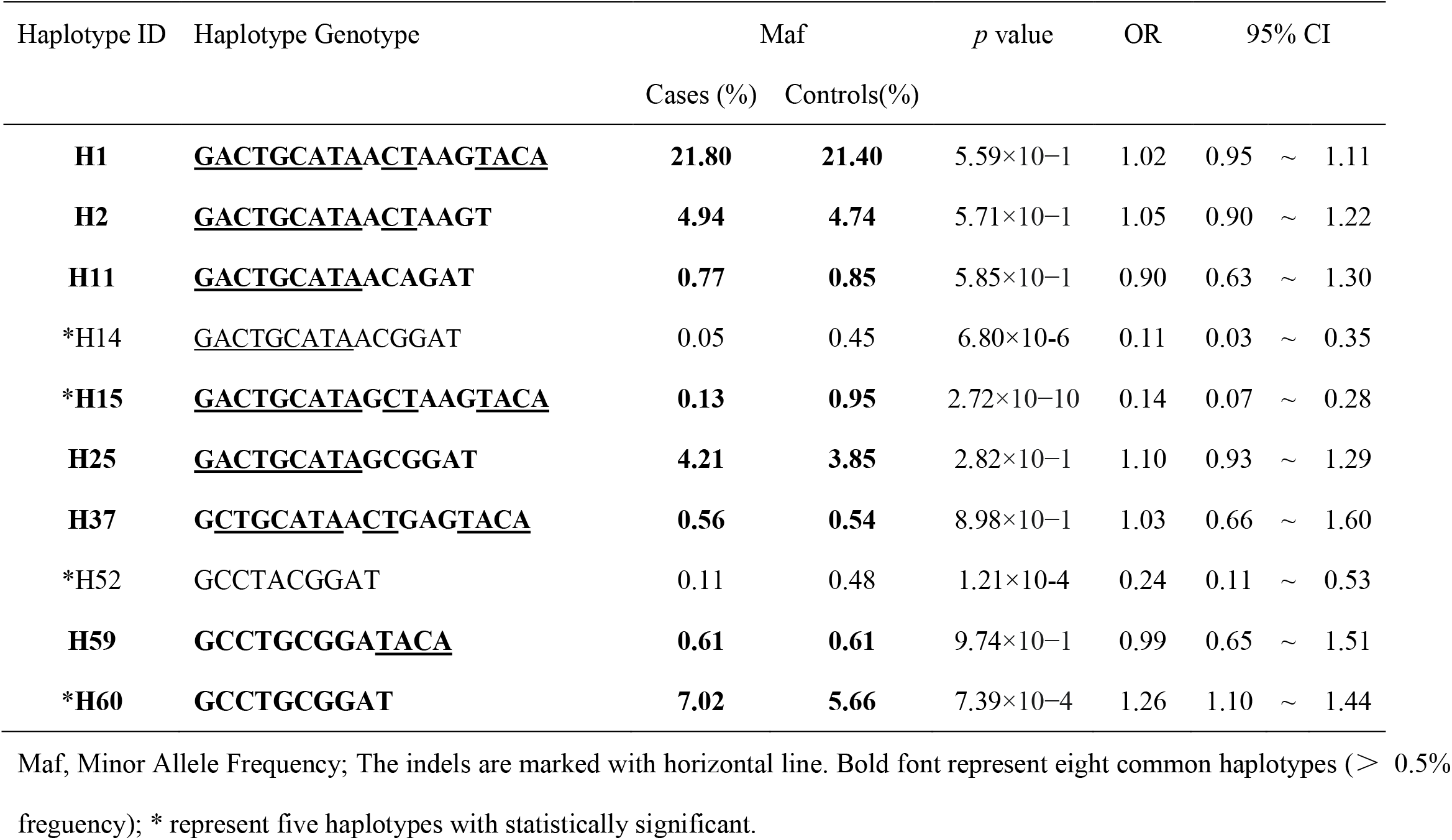
The common and significant haplotypes associated with atopic dermatitis in 5q22.1region

### 3.2. The results of targeted next generation sequencing

Sixteen individuals (eight cases and eight controls) were selected for sequencing, which represented eight common haplotypes (> 0.5% frequency). Eight cases included five with H1/H15, two with H15/H40, and one with H15/H26; eight controls consisted of one with H15 homozygous, five with H15 heterozygous (H15/H37, H15/H25, H11/H15, H15/H60, H1/H15) and two without H15 (H2/H25, H37/H59) (Supplementary Table 2)

We performed sequencing of 16 DNA samples with an average of 1,629.37 Mb raw bases. After removing low-quality reads, we obtained an average of 10,760,164 clean reads (1,602.37 Mb). The average GC content was 40.21%. The 1.79 Mb target region were captured, and 66.46% mapped to target regions of total effective bases. The mean sequencing depth on target regions was 499.02-fold.

The H15 was considered a research point to discover the causal mutation, but no co-segregated LOF variants with H15 were found in sequencing analysis, indicating that there is no gene mutation specific to H15. Since there were 11 genes in 5q22.1 regions included in the target sequence experiment, to explore whether these genes that have more than two LOF variants are associated with AD, we performed a chi-square test between case and control cohort by using LOF variants frequency that was calculated within one specific gene. The result indicated that TMEM232 was statistically significant associated with AD (*P* = 7.33×10 ^−5^, OR = 0.33, 95% CI = 0.19 -0.58) and had the same direction as H15 (Table2).

**Table 2.** The results of gene level loss of function variants enrichment test

| Gene | cases |  | controls |  | chi-square | *P* value | OR | 95% CI |  |  |
| --- | --- | --- | --- | --- | --- | --- | --- | --- | --- | --- |
|  | Alt(N) | Ref(N) | Alt(N) | Ref(N) |  |  |  |  |  |  |
| TMEM232 | 20 | 172 | 50 | 142 | 15.72 | $7.33 \times 10^{-5}$ | 0.33 | 0.19 | ~ | 0.58 |
| CAMK4 | 23 | 73 | 24 | 72 | 0.03 | $8.67 \times 10^{-1}$ | 0.95 | 0.49 | ~ | 1.83 |
| EPB41L4A | 155 | 149 | 157 | 147 | 0.03 | $8.71 \times 10^{-1}$ | 1.00 | 0.71 | ~ | 1.34 |
| LOC102467214 | 47 | 49 | 46 | 50 | 0.02 | $8.85 \times 10^{-1}$ | 1.04 | 0.59 | ~ | 1.84 |
| MAN2A1 | 118 | 90 | 118 | 90 | 0.00 | / | 1.00 | 0.68 | ~ | 1.47 |
| SLC25A46 | 34 | 142 | 45 | 131 | 1.98 | $1.60 \times 10^{-1}$ | 0.70 | 0.42 | ~ | 1.16 |
| NREP | 6 | 106 | 5 | 107 | 0.10 | $7.57 \times 10^{-1}$ | 1.21 | 0.37 | ~ | 4.09 |
| STARD4 | 4 | 28 | 7 | 25 | 0.99 | $3.20 \times 10^{-1}$ | 0.51 | 0.13 | ~ | 1.95 |
| TSLP | 24 | 120 | 20 | 124 | 0.43 | $5.12 \times 10^{-1}$ | 1.24 | 0.65 | ~ | 2.36 |
| WDR36 | 64 | 272 | 67 | 269 | 0.09 | $7.70 \times 10^{-1}$ | 0.95 | 0.65 | ~ | 1.38 |
| LOC100289673 | 0 | 32 | 2 | 30 | 2.07 | $1.51 \times 10^{-1}$ | 1.07 | 0.98 | ~ | 1.17 |

Alt, alternative allele ; Ref, the allele in the reference genome. N, LOF variants counts in case and control cohort

## 4. Discussion

The GWAS studies have identified 34 risk loci for AD, these loci also suggest that genes in immune responses and epidermal skin barrier functions are associated with AD^[15]^. In fact, the overall study of different SNP sites is more conductive for discovering genes associated with a disease or a certain phenotype^[16]^. Theoretically, the SNPs in proximity should exhibit high LD^[17]^, Many studies based on haplotypes can bring us more efficiency than a single SNP study^[18, 19]^.

The haplotypes spanning the 5q22.1 region were constructed by the genotypes of 10 variants from our previous AD studies^[7, 20]^. The frequency of H15 haplotype was the most significantly different between AD patients and controls, so it was considered a research point to discover the causal mutation. By targeted resequencing of risk and non-risk associated haplotypes in the certain locus, the functional risk variants may be identied^[21]^. However, the target sequence experiment failed to identify causal variant that was co-segregated with H15 haplotype in this study.

The LOF variants of certain gene, including of exonic, UTR, splicing site or upstream, have been inconsistently associated with altered gene expression and/or directly with disease^[22]^. In order to find the susceptibility genes associated with AD, the data of target sequence at 5q22.1 region was used to make gene level enrichment analysis. The number of LOF variants at TMEM232 gene was significant associated with AD. It is consistent with our previous result by the fine gene mapping^[7]^. Meanwhile, it was interesting that the associated direction of this gene is exactly the same as H15 haplotype, which indicate there maybe have a linkage between this gene and haplotype H15 while in a large sample size.

TMEM232 belongs to the transmembrane protein family (TMEMs), including TMEM45A, TMEM45B, and TMEM79 and et al, which have been predicted to be components of cellular membranes, such as mitochondrial membranes, the endoplasmic reticulum, lysosomes and the Golgiapparatus^[23]^. A recent study identified nonsense and missense mutations of the TMEM79 gene that encode the protein mattrin in some Irish AD patients who lack an FLG mutation^[24]^. TMEM45Ais associated with the Golgi apparatus, with the trans-Golgi/trans-Golgi network in vitro and in the granular layer in vivo, which shows a strong correlation between TMEM45A expression and epidermal keratinization^[25]^. There are few reports about the TMEM232 gene, and its function is not clear.

There were limitations in this study. Although the target sequence experiment covered 2.9Mb region, it didn’t overlap the entire 5q22 region. Further research on larger sample sizes, including more high frequency and significant haplotypes, is needed to search for functional gene mutations. In addition, more gene function exploration and the possible underlying mechanisms still require further research.

We confirm previous findings that the TMEM232 gene is associated with AD by haplotype analysis and targeted capture sequencing.

## Acknowledgements

We would like to thank the individuals and their families who participated in this project. This study was funded by the Key Project of Natural Science Research in Colleges and Universities in Anhui Province (No. KJ2016A367), and the National Natural Science Foundation of China (No. 81172838, 81972926).

